# The SinR•SlrR heteromer attenuates transcription of a long operon of flagellar genes in *Bacillus subtilis*

**DOI:** 10.1101/2025.01.06.631544

**Authors:** Ayushi Mishra, Abby Jackson, Xindan Wang, Daniel B. Kearns

## Abstract

During growth, *Bacillus subtilis* differentiates into subpopulations of motile individuals and non-motile chains, associated with dispersal and biofilm formation respectively. The two cell types are dictated by the activity of the alternative sigma factor SigD encoded as the penultimate gene of the 27 kb long *fla/che* flagellar operon. The frequency of SigD-ON motile cells is increased by the heteromeric transcription factor SwrA•DegU that activates the *fla/che* promoter. Conversely, the frequency of motile cells is decreased by the heteromeric transcription factor SinR•SlrR, but the mechanism and location of inhibition is poorly understood. Here, using ChIP-Seq analysis, we determine the binding sites of the SinR•SlrR heteromer on the genome. We identified two sites within the *fla/che* operon that were both necessary and sufficient to attenuate transcript abundance by causing premature termination upstream of the gene that encodes SigD. Thus, cell motility and the transition to biofilm formation depend on the expression of a long operon governed by two opposing heteromeric transcription factors that operate at two different stages of the transcription cycle. More broadly, our study serves as a model for transcription factors that control transcriptional elongation and the regulation of long operons in bacteria.

## INTRODUCTION

*Bacillus subtilis* grows as a mixed population of two different cell types; some cells are motile and grow as individuals, while others are non-motile and grow in long chains (Grant 1969; Nishihara 1975; Kearns 2005). The two subpopulations are differentiated by the level and activity of the alternative sigma factor SigD (Marquez 1990; Kearns 2005; Chen 2009; Cozy 2010). Motile cells have high levels of SigD protein and express a regulon containing late flagellar structural proteins and peptidoglycan (PG) lyases that separate cells after division (SigD^ON^ cells) (Serizawa 2004; Kearns 2005; Cozy 2010; Cozy 2012). Conversely, chaining cells have low levels of SigD protein and fail to express both the flagellar proteins and PG lyases, such that cells fail to separate from one another after division (SigD^OFF^ cells) (Marquez 1990; Chen 2009; Cozy 2010). The two cell types are the product of a developmental, epigenetically-inherited switch (Cozy 2010; Norman 2013; Baker 2016), and likely evolved as a “bet-hedging” strategy to compensate for the fact that the assembly of functional flagella takes two to three generations at high growth rates (Guttenplan 2013).

A number of proteins regulation regulate motile cell development. SigD is important; as without it, the SigD-regulon is inactivated and all cells in the population grow as non-motile chains (Helmann 1988; Serizawa 2004; Kearns 2005; Chen 2009). The gene encoding SigD, *sigD*, is found near the 3’ end of the 32-gene, 27-kb long *fla/che* operon that also encodes proteins required early in flagellar biosynthesis (Helmann 1988, Marquez 1994; West 2000; Cozy 2010). Mutation of any of the flagellar structural genes in the *fla/che* operon inactivates the SigD regulon, as cells fail to export the anti-sigma factor FlgM, FlgM accumulates in the cytoplasm, and SigD activity is inhibited (Mirel 1994; Caramori 1996; Frederick 1996; Bertero 1999, Calvo 2015). The promoter of the *fla/che* operon is activated by a heteromeric complex comprised of the response regulator DegU and small protein SwrA, to increase SigD levels and increase the frequency of motile SigD^ON^ cells (Grant 1969; Kearns 2005; Calvio 2005; Tsukuhara 2008; Cozy 2010;Ogura 2012; Mordini 2013; Mishra 2023). Finally, another heteromeric complex of two paralogous DNA binding proteins, SinR and SlrR, has the opposite effect and decreases the frequency of the motile SigD^ON^ state.

SinR is tetrameric phage-like DNA binding protein that binds to and represses the expression of operons that promote biofilm formation (Lewis 1998; Scott 1999; Colledge 2011; Milton 2020). One target of SinR is the *eps* promoter that drives expression of 15 gene products including EpsH, an enzyme involved in the synthesis of the biofilm extracellular polysaccharide (EPS), and EpsE, a bifunctional EPS synthase and inhibitor of flagellar rotation (Kearns 2005; Blair 2008; Guttenplan 2010). At the same location upstream of *eps*, SinR also represses the oppositely-oriented *slrR* gene that encodes SlrR, a SinR-paralog (Chu 2008; Kobayashi 2008) (**Fig 1A**). When SinR binding to DNA is antagonized by even two-fold higher levels of its antagonist SlrA (Kobayashi 2008; Chai 2009; Cozy 2012), SlrR is de-repressed and forms a heteromer with SinR (Chai 2010; Newman 2013). SlrR is not known to bind DNA on its own but is thought to reprogram the binding of SinR to other sites in the chromosome, including those responsible for the inhibition of SigD-dependent gene expression and population heterogeneity (Chai 2010; Cozy 2012). Antagonism of SinR and the formation of the SinR•SlrR heteromer is central to the transition from motility to biofilm formation. How the heterodimer inhibits SigD-dependent gene expression is poorly understood and there are two models for how inhibition might occur.

**Figure 1:**
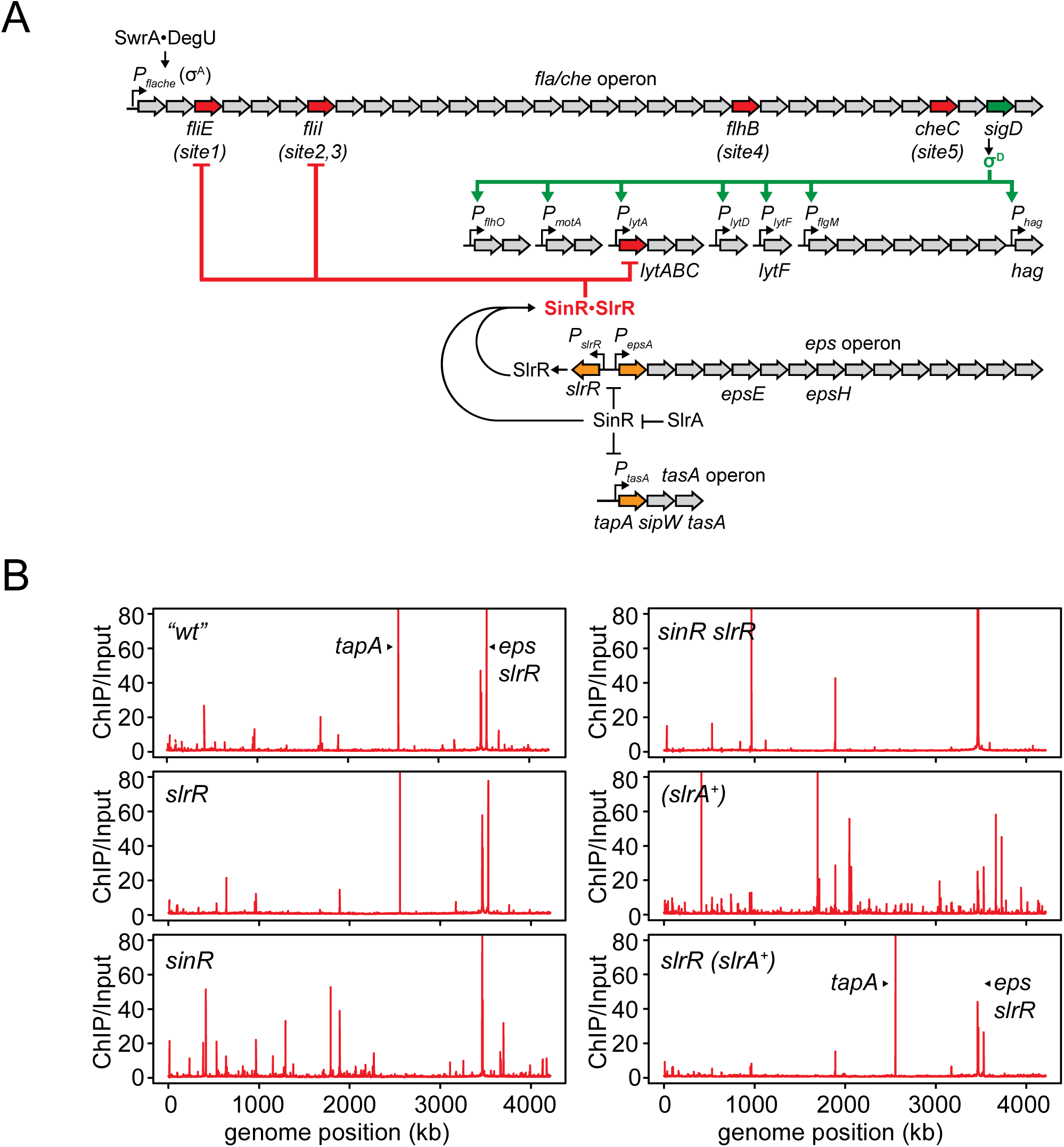
ChIP-Seq analysis indicates different enrichment profiles for SinR, SlrR and the SinR•SlrR heteromer *in vivo*. **A)** A model for SinR•SlrR-mediated inhibition of SigD gene expression. A cartoon diagram of the flagellar and biofilm regulons is presented in the figure. SinR represses the promoter of the *tasA* operon that encodes for protein components of biofilm formation and the promoter of the *eps* operon that encodes genes for the synthesis of the biofilm extracellular polysaccharide (EPS). SinR simultaneously inhibits a divergent promoter that expresses the paralog, SlrR. Genes directly downstream to the promoters inhibited by SinR are highlighted in orange. Elevated levels of SlrA antagonize SinR, promoting formation of the SinR•SlrR heteromer. The heteromer binds to multiple sites within the *fla/che* operon, reducing flagellar gene expression and the levels of SigD, causing a failure in SigD-dependent gene expression. Inhibition by the heterodimer occurs in a subpopulation of cells and can be overridden by SwrA•DegU-dependent activation of the *P_fla/che_* promoter. The Bent arrows indicate promoters. Block arrows indicate genes, and those containing binding sites for SinR•SlrR are highlighted in red. Other arrows indicate activation and T-bars indicate inhibition. The names of relevant genes mentioned in the text are indicated below their corresponding gene location. **B)** ChIP-Seq was performed using a primary antibody to SinR that also cross-reacts with SlrR (Cozy, 2012). For each sample, the number of normalized reads from samples treated with α-SinR was divided by the normalized reads for the corresponding untreated samples to determine fold enrichment (ChIP/Input). ChIP/input reads were plotted in 1 kb bins against genome position in kilobases. Each strain used in this experiment was deleted for *epsH* to abolish cell-clumping that occurs in the absence of SinR. Therefore, an *epsH* mutant was considered “*wt*” for this experiment. The genotype of the strain used to generate each panel is indicated in the top left corner. The following strains were used to generate the data in the indicated panel: “*wt*” (DS6776), *slrR* (DK9313), *sinR* (DK9090), *sinR slrR* (DK9314), (*slrA^+^)* (DK9093), and *slrR (slrA^+^)* (DK9332). The genotype (*slrA^+^)* indicates a “*wt*” strain with an additional copy of *P_slrA_-slrA* integrated at an ectopic locus. Two carets indicate peaks that correspond to the promoter regions of the *eps* operon and the *tapA-sipW-tasA* operon and labelled with the first gene downstream to the promoter region, *epsA* and *tapA* respectively.

One model for SinR•SlrR inhibition of motility gene expression is direct, in which the heteromer inhibits gene expression directly at individual promoters recognized by RNA polyermase and SigD. In support of this model, the heteromer was shown to bind to promoter regions of multiple SigD-dependent genes *in vitro* including those that code for the flagellin protein Hag, and the peptidoglycan lyases LytF and LytABC (Chai 2010). Thus, the heteromer directly blocks expression of motility and cell separation genes at their individual promoters. The “direct inhibition model”, however, cannot explain why SigD fails to accumulate in the presence of the heteromer, or explain the observation that seemingly all of the SigD regulon is repressed, unless the heteromer binds every SigD-controlled promoter (Cozy 2010; Cozy 2012). Another model for SinR•SlrR inhibition of motility gene expression is indirect, in which the heteromer inhibits the expression of SigD itself. In support of this model, the heteromer inhibits accumulation of SigD by inhibiting transcript levels of the *fla/che* operon somewhere between the *P_fla/che_* promoter and the *sigD* gene (Cozy 2012). The “indirect inhibition model”, however, cannot explain where the heteromer might bind within the *fla/che* operon, and even if it did, how transcriptional inhibition might occur. The sequence to which the SinR•SlrR heteromer binds is unknown.

Here we explore the mechanism of SigD inhibition by determining where in the genome SinR, SlrR, and the SinR•SlrR heteromer bind, using chromatin immunoprecipitation and DNA sequencing (ChIP-Seq). Our ChIP-Seq data support known SinR binding sites and reveal that SlrR binds to DNA in the absence of SinR. The data also support the idea that SinR and SlrR form a heteromer that binds to sites different from either homomer alone, with a predicted consensus sequence that resembles directly adjacent half-sites for each protein. Many of the heteromer binding sites were within open reading frames, including five locations within the *fla/che* operon. Genetic analysis indicated that two of the heteromer binding sites within *fla/che* were responsible for decreasing the frequency of SigD-dependent gene expression both when SlrA was expressed in extra copy and when SwrA was absent. Moreover when bound, SinR•SlrR inhibited transcript abundance both at and downstream of the binding sites, likely by impeding elongation and/or promoting premature termination, and each site was sufficient for doing so when integrated in artificial reporter systems. In sum, our data support the indirect inhibition model of motile cell development in *B. subtilis,* in which the SinR•SlrR heteromer inhibits SigD levels. Impairment of transcription elongation by transcription factor binding is considered rare in bacteria, but may be more common than appreciated, and long operons may be particularly susceptible to such attenuation.

## RESULTS

### Homomers and heteromers of SinR and SlrR bind to different sites

To explore the relative regulatory contributions of the paralogs SinR and SlrR, we performed ChIP-Seq analysis on exponentially growing wild type cells using a polyclonal antibody that reacts to both proteins (Cozy, 2012). SinR represses the expression of SlrR during growth and thus any peaks obtained were expected to be largely due to the SinR homomer (**Fig 1A**). Consistent with expectations, relatively few peaks were observed, two of which represented known SinR binding sites, located in the intergenic regions upstream of the gene that encodes SlrR, the *eps* operon, and *tasA* operon (Kearns 2005, Chu 2006) (**Fig 1B**, **Fig 2A**). Also consistent with expectation, mutation of SlrR produced a peak pattern very similar to that observed for wild type (**Fig 1B**), while mutation of SinR abolished the peaks upstream of the *eps* and *tasA* operons (**Fig 1B**, **Fig 2A**, **Table S1**). MEME analysis of 200 base pair sequences surrounding the peak centers of SinR-dependent peaks identified binding sites for SinR, which were similar to the consensus half-site of GTTCTYT that was previously-identified (Chu 2006) (**Fig 2B, Table S1**). We conclude that our genome-wide analysis supports pre-existing data and models where SinR is the predominant repressor that binds to and represses both SlrR and the loci required for biofilm formation.

**Figure 2:**
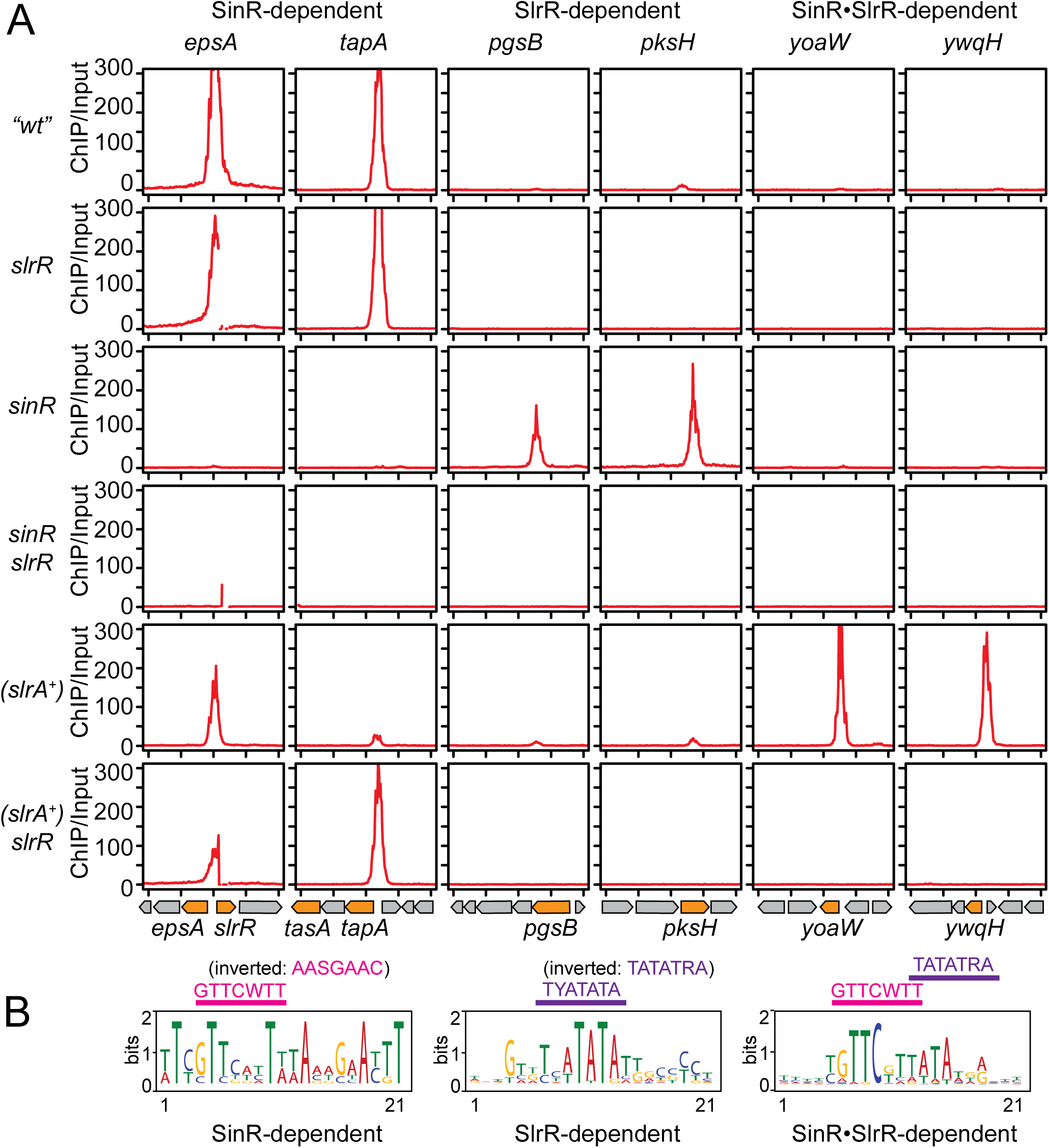
MEME analysis indicates distinct sequence patterns for SinR homomer, SlrR homomer and SinR•SlrR heteromer-dependent enrichment. **A)** Enlarged view of regions of interest from data presented in Figure 1B. Analysis was performed the same way as in Figure 1B and the ChIP/input (Y-axis) was plotted in 10 bp bins over a 4 kb range. Ticks on the X-axis represent 1 kb intervals. The neighboring genome architecture is cartooned below each vertical panel and relevant genes of interest are colored orange. Each vertical panel is labelled after the gene or promoter region within which the peak was found. The following strains were used to generate this figure. “*wt*” (DS6776), *slrR* (DK9313), *sinR* (DK9090), *sinR slrR* (DK9314), (*slrA^+^)* (DK9093) and *slrR (slrA^+^)* (DK9332). Note, a discontinuity is observed in some of the peaks in the left-most column due to deletion of *slrR*. **B)** A 200 bp region around the center of peaks in each category was subjected to MEME analysis and a 21 bp MEME was generated. The predicted SinR half-site is marked in pink and is consistent with reported binding sites (Kearns, 2005). The predicted half-site of SlrR is marked in purple. The following abbreviations were used to describe base conservation: W (A or T), S (G or C), Y (C or T), and R (A or G).

In the wild type, peaks were enriched by ChIP-Seq that could not be attributed to SinR binding, and new peaks were enriched in a SinR mutant (**Fig 1B**). Some of the peaks in the SinR mutant were possibly due to de-repression of SlrR, but to the best of our knowledge SlrR has not been reported to bind DNA or regulate gene expression on its own. ChIP-Seq analysis of a strain doubly mutated for SinR and SlrR abolished 20 peaks, which were deemed to be SlrR-dependent (**Fig 1B**, **Fig 2A, Table S2**). Unlike the intergenic SinR-dependent peaks, the SlrR-dependent peaks were found to be overwhelmingly located within open reading frames (**Table S2**), and MEME analysis of 200 base pair sequences surrounding the peak centers indicated a putative consensus half-site of TYATATA (**Fig 2B**). Finally, some peaks remained in the absence of both SinR and SlrR and we wondered whether these might be due to one or more of the seventeen other Xre transcription factor family paralogs encoded by *B. subtilis*. Simultaneous mutation of SinR, SlrR, and the paralog YgzD, abolished a single additional peak upstream of the gene that encodes YgzD and the *ygzD* promoter was found to be auto-repressed (**Fig S1, S2**). We conclude that the polyclonal antibodies originally raised against SinR (Kearns 2005), cross-react with SlrR, YgzD and potentially other members of the family. We further conclude that SlrR and other SinR-paralogs bind to specific locations in the chromosome.

Previously reported genetic and biochemical data suggest that SinR and SlrR can form a heteromer that reprograms SinR to bind new sites in the chromosome when the small antagonist protein SlrA is in excess (Chai 2010; Cozy 2012). Extra SlrA is thought to disrupt a subpopulation of SinR homomers, partially de-repress expression of SlrR and facilitate heteromer formation. To test for the binding of the heteromer, ChIP-Seq was performed on a strain that encoded an extra copy of the *slrA* gene expressed from its native promoter and integrated at an ectopic site in the chromosome (*slrA^+^*) (**Fig 1B**). When *slrA* was present in extra-copy, SinR-dependent peaks were diminished, perhaps consistent with partial SinR antagonism, and a number of new peaks that were not previously attributed to either SinR or SlrR alone, were observed (**Fig 1B**, **Fig 2A, Table S3**). As with the putative SlrR homomer, the peak sites for the heteromer were largely intragenic, and mutation of SlrR abolished the additional peaks (**Fig 1B**, **Fig 2A**). MEME analysis of 200 base pair sequences in the centers of the putative heteromer peaks indicated an elongated consensus sequence of GTTCWTTATATRA (**Fig 2B**). We note that the consensus appears to be the half-sites of SinR and SlrR respectively, directly juxtaposed. We conclude that the SinR•SlrR heteromer binds to a new set of genes that are not direct targets of either homomer.

### SinR•SlrR binds within *lytA* and multiple sites within the *fla/che* operon

One model to explain how the SinR•SlrR heteromer inhibits SigD-dependent gene expression is by direct repression of the SigD-dependent genes *lytABC*, *lytF* and *hag* (Chai, 2010). No peaks were detected near *lytF* or *hag* in any of the ChIP-Seq experiments, but a peak was detected near the *lytA* promoter region (**Fig 3A**). MEME analysis suggested a putative heteromer binding site was located within the *lytA* open reading frame near the 5’ end, and mutation of SlrR but not SinR, abolished the peak (**Fig 3A**). To test the role of the putative binding site in the regulation of *lytA*, two promoter fusions were generated to the *lacZ* gene encoding β-galactosidase, one which included just the intergenic region upstream of *lytA* (*P_lytA_-lacZ*) and one which included the intergenic region plus the putative intragenic binding site (*P_lytA_^ext^-lacZ*) (**Fig 4A**). Mutation of SinR or both SinR and SlrR together, did not alter expression of either reporter suggesting that the ChIP-Seq enrichment by SlrR alone was inconsequential (**Fig 4B**). In the presence of an extra copy of the *slrA* gene, however, expression of both reporters was reduced, and was restored when SlrR was also mutated, consistent with heteromer repression (**Fig 4B**). We conclude that the heteromer inhibits expression of the *P_lytA_* promoter but whether the effect is direct or indirect is unclear as it appeared to be independent of the putative binding sequence.

**Figure 3:**
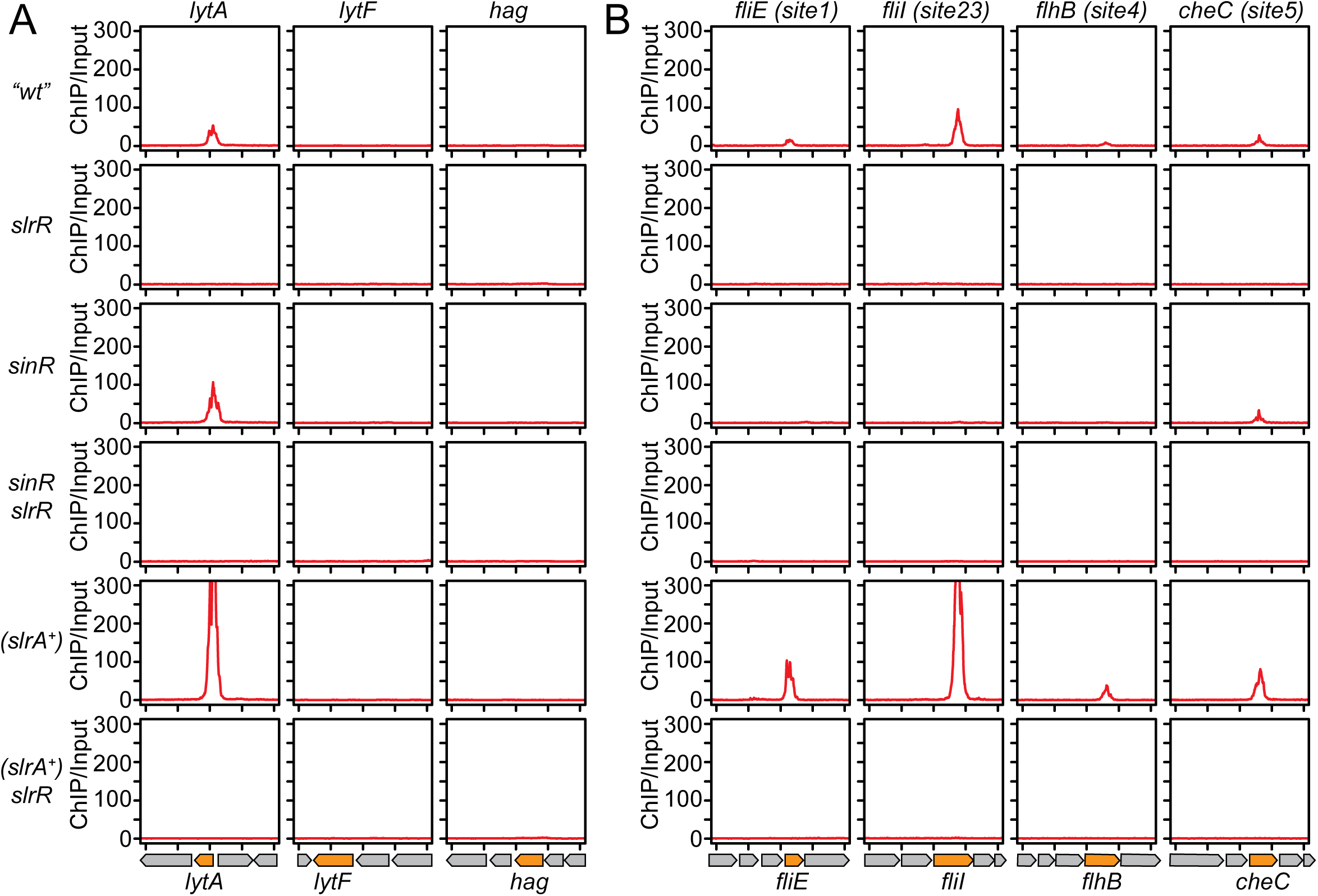
The SinR•SlrR heteromer is enriched at four locations within the *flache* operon. **A)** Enlarged view of regions surrounding previously reported SinR•SlrR heterodimer target genes: *lytA*, *lytF* and *hag*. Analysis was performed the same way as in Figure 1B and the ChIP/input (Y-axis) was plotted in 10 bp bins over a 4 kb range. Ticks on the X-axis represent 1 kb intervals. Each vertical panel corresponds to a particular gene or promoter region indicated at top. The neighboring genome architecture is drawn below each vertical panel and relevant genes of interest are colored orange. **B)** Enlarged view of regions enriched by the heteromer within the *fla/che* operon. A total of five putative binding sites were identified in four locations, and the sites were named *site1* through *site5*. Each vertical panel corresponds to a particular gene indicated at top. The following strains were used to generate this figure. “*wt*” (DS6776), *slrR* (DK9313), *sinR* (DK9090), *sinR slrR* (DK9314), (*slrA^+^)* (DK9093) and *slrR* (*slrA^+^)* (DK9332).

**Figure 4:**
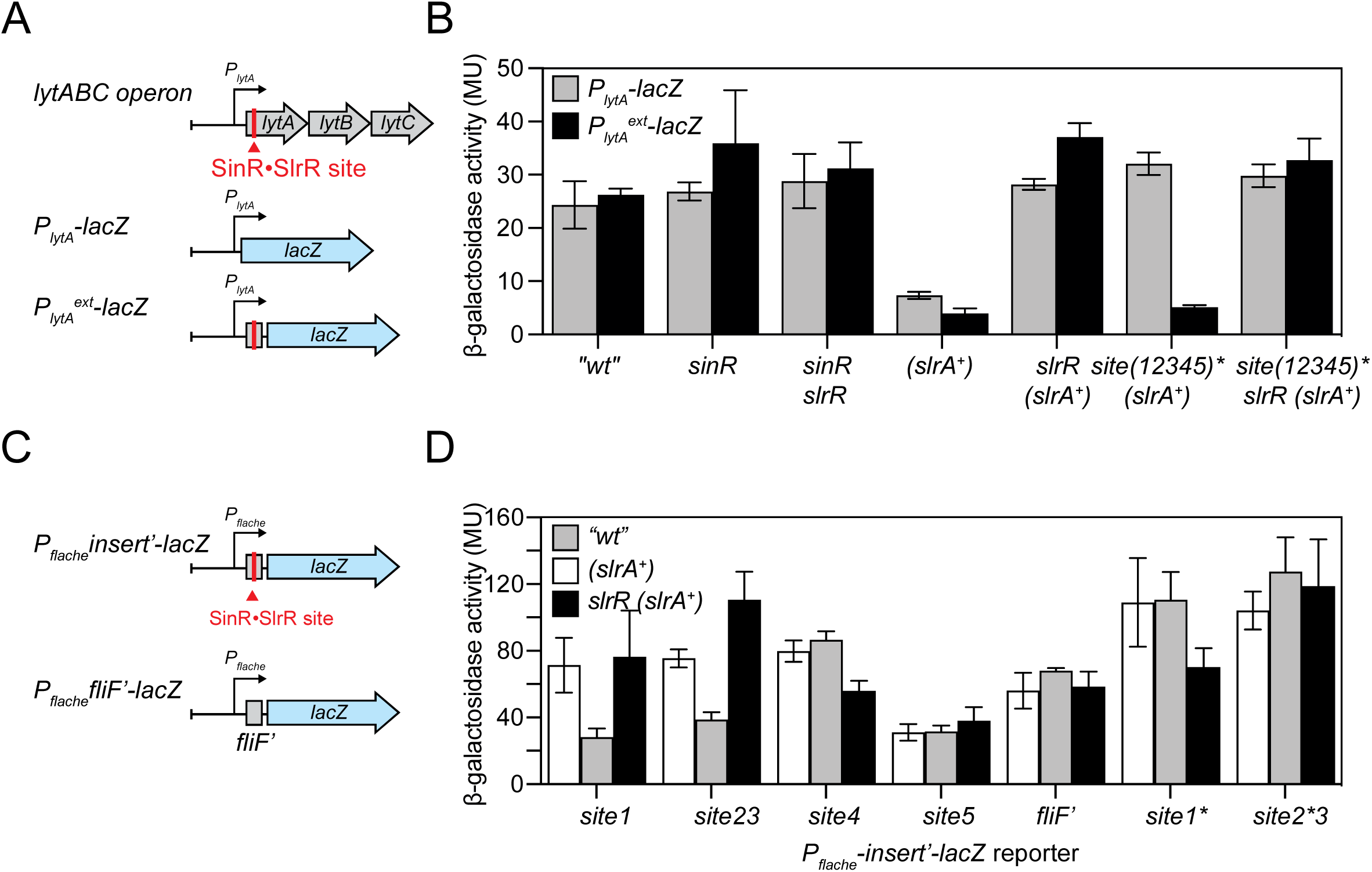
SinR•SlrR binding sites within *fla/che* and *lytA* are both necessary and sufficient for transcription inhibition. **A)** A schematic of *lytA* genomic region and the two reporters, *P_lytA_*-*lacZ* and *P_lytA_^ext^*-*lacZ* that were used in this experiment. *P_lytA_*-*lacZ* includes the promoter region of *lytA* upstream transcriptionally fused to *lacZ*. *P_lytA_^ext^*-*lacZ* includes the upstream region of *lytA* and the predicted SinR•SlrR binding site within *lytA* open reading frame transcriptionally fused to *lacZ*. **B)** β-galactosidase activity in Miller units (MU) plotted on the Y-axis in linear scale. Each experiment was performed in three replicates and error bars indicate standard deviation of the three replicates. Gray bars represent expression of *P_lytA_*-*lacZ* and black bars represents expression of *P_lytA_^ext^*-*lacZ*. The genetic background in which each reporter was tested is indicated on the X-axis. Each strain used in this experiment was deleted for *epsH* to abolish for cell-clumping in the absence of SinR, and thus an *epsH* mutant was considered “*wt*” for this experiment. The following strains were used to generate this panel, “*wt*” (DB1829, DB1834), *sinR* (DB1830, DB1835), *sinR slrR* (DB1831, DB1836), (*slrA^+^)* (DB1832, DB1837), *slrR* (*slrA^+^)* (DB1845, DB1846), *site(12345)\** (*slrA^+^)* (DB1833, DB1838) and *site(12345)* slrR* (*slrA^+^)* (DB1880, DB1881). Raw data are presented in **Table S4**. **C)** A schematic of Transcriptional *lacZ* reporters of *P_flache_* fused to 150 bp region surrounding the SinR•SlrR site within *fliE* (*site1*), *fliI* (*site23*), *flhB* (*site4*) and *cheC* (*site5*) and a site within *fliF* not associated with a heteromer binding peak in ChIP-seq analysis. Similar reporters were also constructed where *P_flache_* was fused to *site1\** and *site2\** mutants respectively. **D)** β-galactosidase activity in Miller units (MU) plotted on the Y-axis in linear scale. Error bars are the standard deviation of three replicates. Each strain used in this panel was mutated for *epsE* to avoid clumping of cells in the absence of SinR and thus, an *epsE* mutant was considered as “*wt*” for this experiment. Each reporter was tested in “*wt*” (white bars), *slrA^+^* (gray bars) and *slrR slrA^+^* (black bars) genetic backgrounds. The following strains were used in this experiment, *site1* (DB1777, DB1765, DB1771), *site23* (DB1778, DB1766, DB1772), *site4* (DB1779, DB1767, DB1773), *site5* (DB1780, DB1768, DB1774), *fliF*’ (DB1781, DB1769, DB1775), *site1\** (DB1888, DB1889, DB1890) and *site2*3* (DB1891, DB1892, DB1893). Raw data are presented in **Table S5**.

Another model to explain how the SinR•SlrR heterodimer inhibits SigD-dependent gene expression is by inhibiting expression within the *fla/che* operon somewhere downstream of the *P_flache_* promoter and upstream of the gene encoding SigD. Consistent with the previous work suggesting that the *P_flache_* promoter was not a direct target, no peaks were detected near the *P_flache_* promoter in any of the ChIP-Seq experiments (Cozy 2012). Instead, four peaks were observed within the *fla/che* operon, and MEME analysis indicated five heteromer binding sites centered within the ChIP-Seq peaks. Thus, we named the putative sites: *site1* (within *fliE*), *site2* and *site3* (within *fliI*), *site4* (within *flhB*) and *site5* (within *cheC*). One way in which the heteromer could impair *sigD* expression is by binding sites to the sites and repressing internal promoters. No β-galactosidase activity was detected, however, when a 500 bp region encompassing the peak center for each site was separately cloned upstream of the *lacZ* gene and inserted at an ectopic site in the wild type chromosome (**Fig S3**). We conclude that if the putative heteromer binding sites inhibit *sigD* gene expression, they do not do so by repressing nearby promoters.

To test the role of *sites 1* through *5*, silent mutations were individually introduced in each of the SinR consensus half-sites such that the DNA binding sequence was altered but the protein code was not (**Fig S4A**). None of the silent mutations impaired swarming motility of the wild type, suggesting each was neutral on the effect of their respective flagellar genes (**Fig S4B**). Next, an extra copy of *slrA* was introduced to assess the effect of the heteromer on motility, and while mutation of *site1* caused a slight increase, none of the single point mutations was sufficient to restore motility to wild type levels (**Fig S4B**). Finally, reporters for SigD-dependent gene expression in which the promoter of the *hag* flagellin gene was cloned upstream of either the *lacZ* gene or the *gfp* gene encoding green fluorescent protein (GFP) were introduced (**Fig 5, S5**). Whereas wild type cells expressed high levels of β-galactosidase and GFP fluorescence, cells containing an extra copy of *slrA* were strongly inhibited for both reporters (**Fig 5**). Moreover, expression was restored to both reporters when SlrR was mutated, but none of the single site mutations were sufficient to restore wild type levels of LacZ or GFP expression. We conclude that none of the binding site mutations were sufficient to restore wild swarming motility or SigD-dependent gene expression when inhibited by the SinR•SlrR heteromer. We note however, that minor rescue phenotypes may indicate additive effects of multiple sites within the operon.

**Figure 5:**
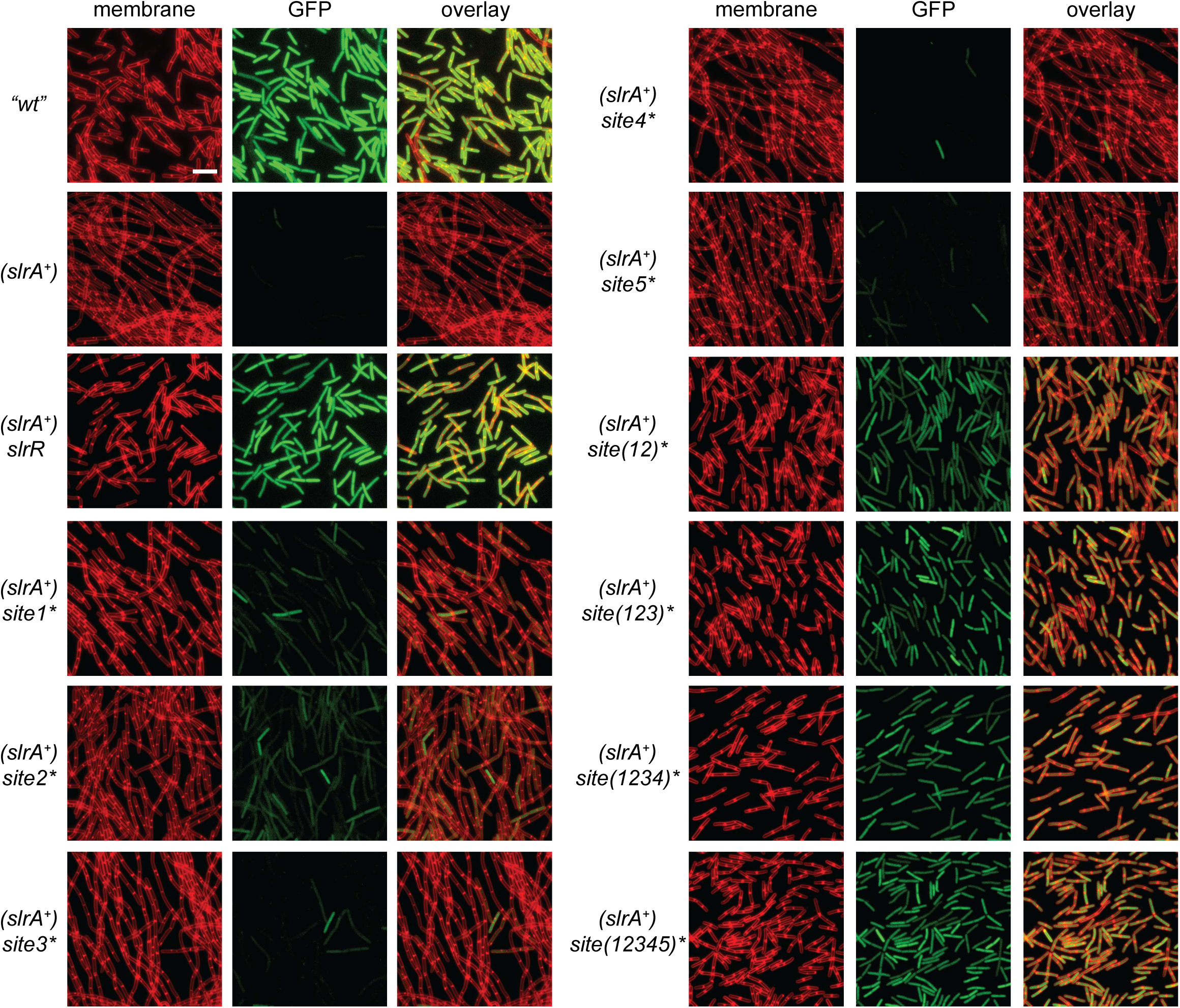
Mutation of SinR•SlrR sites 1 and 2 together is necessary to restore *P_hag_*-GFP expression and cell separation in cells expressing an extra copy of *slrA*. Fluorescent micrographs of cells that contain a *P_hag_-GFP* reporter for SigD-dependent gene expression (GFP, false colored green) and stained with FM 4-64 (membrane, false colored red). All strains used in this panel are mutated for *epsE* (“*wt*”) and thus an *epsE* mutant was considered as “*wt*” for this experiment. The following strains were used to generate the panels: “*wt*” (DB457), (*slrA^+^)* (DB1306), *slrR* (*slrA^+^)* (DB1305), *site1\** (*slrA^+^)* (DB1449), *site2\** (*slrA^+^)* (DB1450), *site3\** (*slrA^+^)* (DB1451), *site4\** (*slrA^+^)* (DB1447), *site5\** (*slrA^+^)* (DB1448), *site(12)\** (*slrA^+^)* ( DB1549), *site(123)\** (*slrA^+^)* (DB1680), *site(1234)\** (*slrA^+^)* ( DB1675), *site(12345)\** (*slrA^+^)* (DB1710). Scale bar is 8μm.

### Multiple SinR•SlrR binding sites within the *fla/che* operon are necessary for inhibiting SigD-dependent gene expression

To test the possibility that each of the putative SinR•SlrR binding sites within the *fla/che* operon had an additive effect on SigD inhibition, mutations were sequentially added until all five sites had been disrupted. Simultaneous mutation of *site1\** and *site2\** (*site(12)*\*) restored partial swarming motility when an extra copy of *slrA* was present, and motility was further improved by additional mutations such that the quintuple mutant exhibited wild type swarming rates, albeit with an extended lag period (**Fig S4B**). Likewise, mutation of *site(12)*\* in cells containing an extra copy of *slrA* increased *P_hag_* expression to near wild type levels, but mutation of additional sites did little to improve expression further (**Fig 5, S5**). Finally, ChIP-Seq analysis in the quintuple mutant in cells containing extra an extra copy of *slrA* indicated that mutation of first four sites abolished enrichment at their respective locations but mutation of *site5\** did not (**Fig S6**). We conclude that *site1* and *site2* are required for SinR•SlrR binding *in vivo* and play a predominant role in inhibiting both swarming motility and SigD-dependent gene expression.

To determine if any of the SinR•SlrR predicted sites within the *fla/che* operon were sufficient to inhibit transcription, reporters were generated in which 200 base pairs surrounding the peaks corresponding to *site1*, *site23*, *site4*, *site5*, and a control fragment from the gene *fliF,* were cloned between the *P_flache_* promoter and the *lacZ* gene (**Fig 4A**). Expression from the reporters with an intervening sequence containing either *site1* or *site23* were reduced in expression when *slrA* was present in extra copy, but the remaining reporters were unaffected (**Fig 4B**). Moreover, the expression levels of both the *site1* and *site2* containing reporters was restored either in the absence of SlrR or when mutations that altered the putative binding site (*site1\** and *site2\**, respectively) were introduced into the intervening sequence (**Fig 4B**). We conclude that while not all heteromer binding sites are necessarily relevant, the *site1* and *site2* cis-elements specifically attenuate transcription shen cloned between a promoter and reporter gene.

To directly observe the effect of *site1* and *site2* on transcript abundance of the *fla/che* operon, RNAseq was performed in wild type and a variety of mutants. Transcript per million (TPM) values of each gene was calculated and normalized to the TPM of *sigA*, gene encoding the housekeeping sigma factor SigA. Similar to a previous report (Cozy, 2012), transcript levels of the *fla/che* operon were high in the *wild type*, but decreased in abundance in when *slrA* was provided in extra copy, specifically near *site1*, with a further decrease that was observed near *site2* (**Figure 6**). Mutation of *site1\** in the strain containing an extra copy of *slrA* raised transcript levels to that of wild type early in the operon, but transcript levels decreased near *site 2* and persisted at a low level (**Figure 6**). Finally, either mutation of both *site1* and *site2* simultaneously (*site(12)*\*), or mutation of SlrR restored transcript to wild type levels to the *fla/che* operon and SigD-regulon in the presence of an extra copy of *slrA* (**Figure 6, Figure S7**). We conclude that heteromer binding to either *site1* or *site2* is both necessary and sufficient to attenuate transcript abundance downstream of promoter initiation. We further conclude that transcriptional attenuation within the *fla/che* operon contributes to the reduction in SigD protein levels to impair expression of the SigD-regulon. We note that of the genes we examined in the SigD regulon, transcript abundance of *lytA* was unusual in that it was restored by mutation of SlrR but not the *site(12)\** mutation (**Figure S7**).

**Figure 6:**
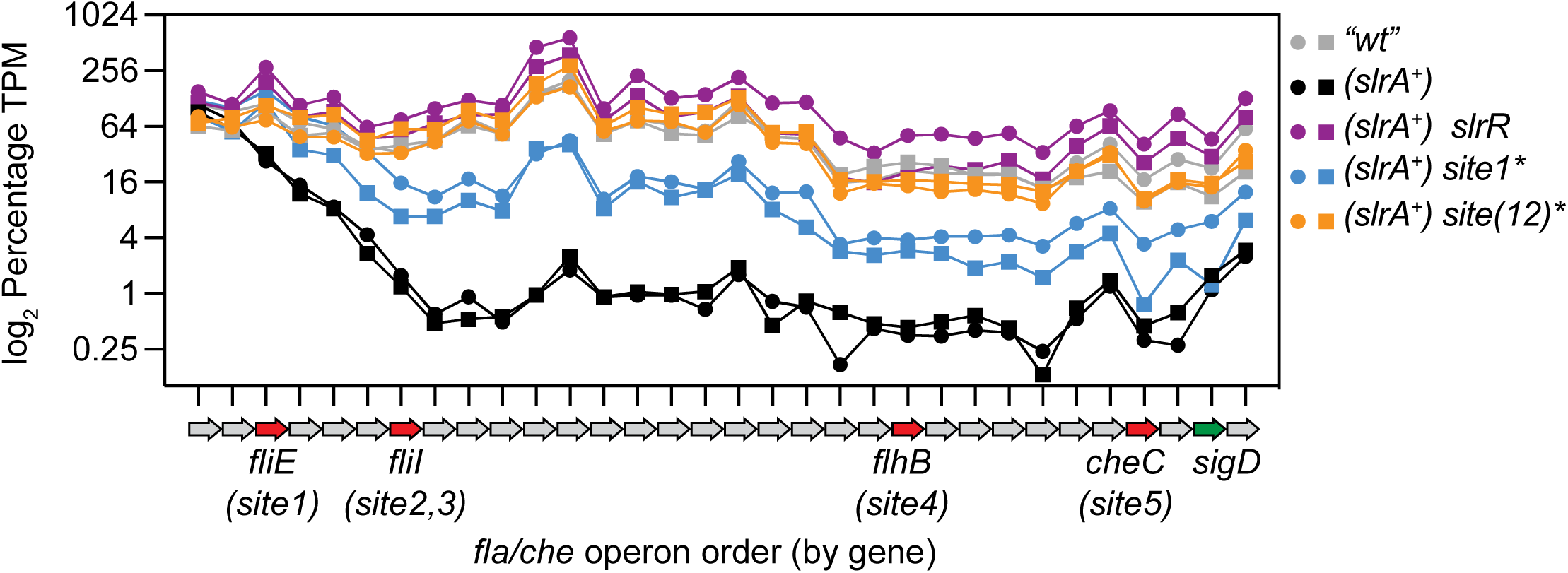
Binding of the SinR•SlrR heterodimer reduces transcript abundance within the *fla/che* operon. RNAseq transcriptomics were performed in duplicate (circles and squares) on the following strains. All strains were mutated for *epsE* (“*wt*”) for consistency with other experiements. “*wt*” (grey symbols, DK9699), (*slrA^+^)* (black symbols, DB154), (*slrA^+^) slrR* (purple symbols, DB141), (*slrA^+^) site1\** (blue symbols, DB1438), and (*slrA^+^) site(12)\** (orange symbols, DB1522). Transcript abundance is expressed by gene position in the *fla/che* operon from 5’ to 3’ end (X-axis) as log_2_ of the percentage transcript relative to the control transcript for the vegetative sigma factor SigA (Y-axis). Raw data are available in **Table S10**.

As the SinR•SlrR heteromer inhibits SigD accumulation, and expression from *P_lytA_* is SigD-dependent, we wanted to determine why transcript abundance of *lytA* failed to increase when SigD activity was restored by mutation of the heteromer binding sites within *fla/che*. Simultaneous mutation of *sites(12345)*\* in the *fla/che* operon, restored expression of *P_lytA_-lacZ* when an extra copy of *slrA* was present, consistent with the indirect model in which the heteromer acts through inhibiting SigD levels, but inconsistent with the low level of *lytA* transcript expression observed by RNAseq (**Figure 4B, Figure S7**). Mutation of *sites(12345)\** however, did not restore expression to the reporter that included the intragenic binding site (*P_lytA_^ext^-lacZ*) in the presence of an extra copy of *slrA*, thereby supporting a model where the heterodimer also directly inhibits *lytA* (**Fig 4B**). Finally, mutation of SlrR restored wild type levels of expression to the extended reporter when *slrA* was in extra copy and the *fla/che* operon binding sites were mutated (**Fig 4B**). We conclude that the heteromer primarily represses the SigD regulon by binding within the *fla/che* operon, but heteromer binding can also have direct effects within individual target genes. Thus, both models for transcriptional inhibition of the SigD regulon are at work, at least for *lytA* gene.

To promote the formation of the SinR•SlrR heteromer, a strain that expresses an extra copy of the *slrA* gene has been used that switches the population heavily in favor of the SigD-OFF state. Population heterogeneity with respect to motility, however, was first observed as both a reduction in fluorescence intensity and a reduction in the frequency of fluorescent SigD-ON cells in a strain mutated for the flagellar master activator protein SwrA and containing a *P_hag_*-GFP reporter (Kearns 2005) (**Fig 7**). Previous work indicated that the SinR•SlrR heteromer played a role as mutation of SlrR increased the frequency of SigD-ON cells in the absence of SwrA (Chai 2010; Cozy 2012) (**Fig 7**). Here we find that mutation of both *site1\** and *site2\** simultaneously, but not either site alone, was sufficient to increase the frequency of SigD-dependent gene expression in cells lacking SwrA, thereby phenocopying the absence of SlrR (**Fig 7**). Finally, the magnitude of SigD-dependent gene expression was not increased in either the SlrR mutant or the *site(12)\** double mutant, consistent with the heteromer acting downstream of SwrA activation at the *P_flache_* promoter (Kearns 2005; Tsukahara 2008; Mordini 2013; Mishra 2024). We conclude that the SwrA•DegU heteromer increases the magnitude of *fla/che* operon expression at the level of transcript initiation, while the SinR•SlrR heteromer attenuates transcript abundance within the operon. Together the two heteromeric systems calibrate the frequency at which the SigD-regulon is activated.

**Figure 7:**
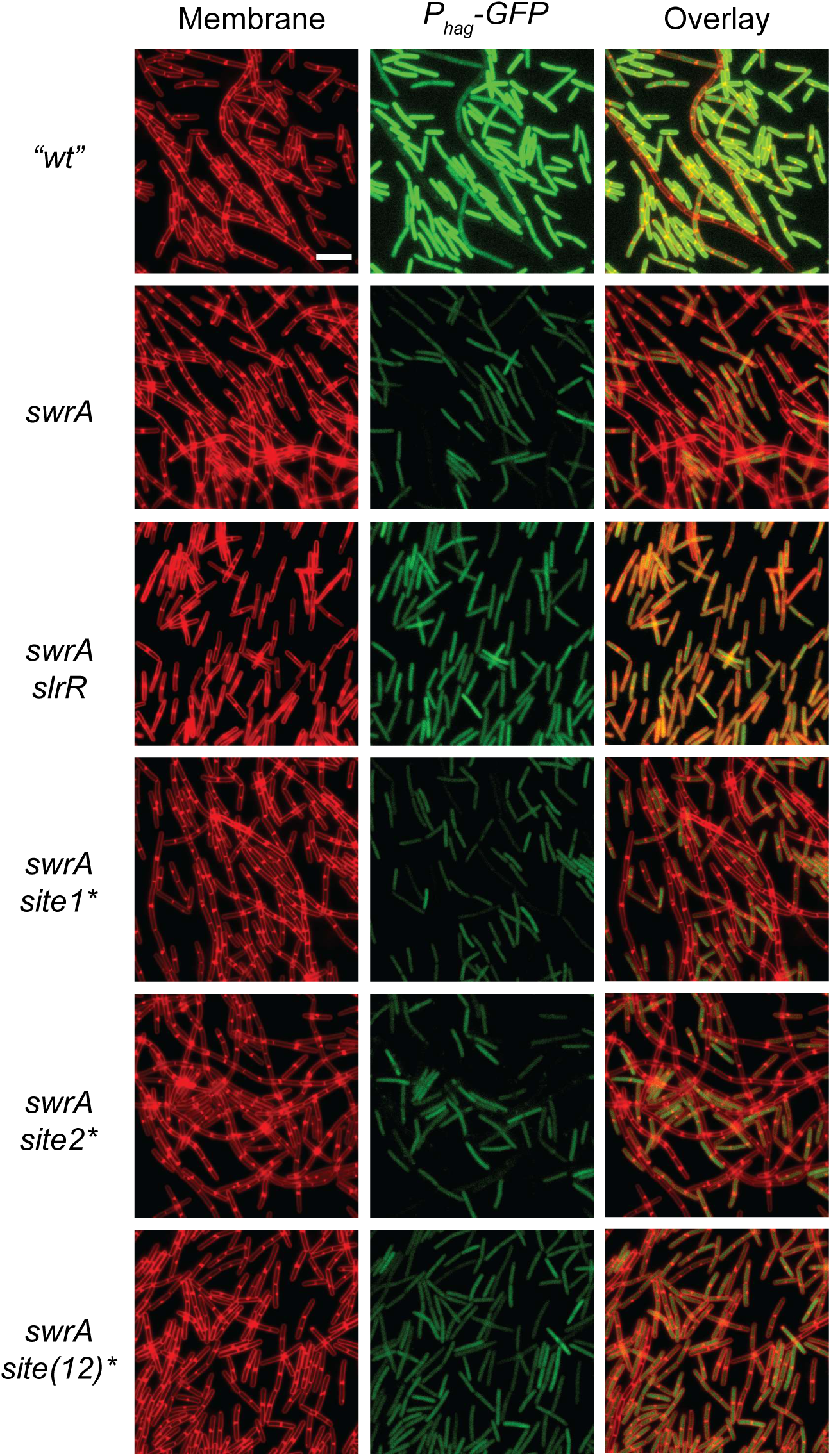
Double mutation of SinR•SlrR *sites(12)\** restores frequency of *P_hag_*-GFP expression in cells lacking SwrA. Fluorescent micrographs of cells that contain a *P_hag_-GFP* reporter for SigD-dependent gene expression (GFP, false colored green) and stained with FM 4-64 (membrane, false colored red). All strains used in this panel are mutated for *epsE* and an *epsE* mutant was considered as “*wt*” for this experiment to maintain consistency with other figures in the manuscript. *P_hag_*-GFP expression in the indicated genetic background is represented in the fluorescent micrographs. *“wt”* (DB457), *swrA* (DB1456), *swrA slrR* (DB1457), *swrA site1\** (DB1444), *swrA site2\** (DB1445), and *swrA site(12)\** (DB1543). Scale bar is 8μm.

## Discussion

Bacteria were once thought to be physiologically uniform during exponential growth, but growing *B. subtilis* spontaneously bifurcates into two phenotypically-distinct subpopulations: single motile cells and long non-motile chains (Kearns and Losick, 2005). Each cell type is differentiated at the level of gene expression governed in part, by the phage-like DNA binding protein SinR (Chai 2010, Cozy 2012). SinR represses genes involved biofilm formation as a homomer (Kearns 2005, Chu 2006, Chu 2008), but transient antagonism relieves repression of a paralog called SlrR to form a SinR•SlrR heteromer (Kobayashi 2008; Chai 2009). Genetic evidence indicates that the heteromer reprograms SinR to bind to new sites in the genome that repress a regulon for flagellar assembly and cell separation under the control of the alternative sigma factor SigD (Chai 2010, Cozy 2012). Here we show that the heteromer binds to multiple sites within the long *fla/che* operon that are both necessary and sufficient for attenuating transcript abundance, likely by promoting RNA polymerase pausing and premature termination. The gene encoding SigD is downstream of the binding sites, and we conclude that the SinR•SlrR heteromer indirectly inhibits SigD activity by preventing SigD accumulation above a threshold in a subpopulation of cells (Cozy 2010; Cozy 2012) (**Fig 1A**).

Repressors commonly inhibit transcriptional activation by binding to sites that occlude access of RNA polymerase to promoter elements (Swint-Kruse 2009; Lewis and Adhya 2015; Browning 2019). Sometimes, repressors bind within open reading frames near the promoter and alter promoter access remotely by DNA looping or other changes in conformation. DNA binding repressors that bind within genes to inhibit elongation/promote premature termination like the SinR•SlrR heteromer are rare but one example is the global regulator CodY of *B. subtilis* (Sonenshein 2007). CodY represses transcriptional initiation of many genes in response to cellular GTP and amino acids levels (Ratnayake-Lecamwasam 2001; Brinsmade 2010; Brinsmade 2014), but like SinR•SlrR, CodY also binds within the *fla/che* operon to antagonize SigD, perhaps as a “roadblock” to transcriptional elongation (Belitsky 2011; Ababneh 2015). How DNA binding proteins would inhibit RNA polymerase transcription bubble progression, however, is unclear as often, a dimer binds on the same surface of the dsDNA making impairment of unwinding unlikely. We note however that the SinR and SlrR inverted half-sites directly abut in the case of heteromer binding, spanning roughly 14 nucleotides. As 10 nucleotides constitute a helical turn, we speculate that binding of the heteromer might wrap all the way around the DNA and act as a clamp. While some RNA readthrough was observed at heteromer bound sites, the binding might pause RNA polymerase long enough to promote premature termination and cause polarity on downstream gene expression.

By whatever mechanism heteromer binding inhibits transcription, we note that not all binding sites were effective. For example, while *site1* and *site2* within the *fla/che* operon both induced a local decrease in transcript abundance at both the native site and in heterologous reporter assays, but *site3*, *site4*, and *site5* did not. Moreover, ChIP-Seq analysis indicated that SlrR alone could bind DNA within open-reading frames, but seemed unable to inhibit transcription. In the case of the *lytA* gene, both SlrR-dependent heteromer and SlrR-dependent homomer enrichment was substantial, but the effect of transcription inhibition was only observed when SinR was present. Thus, we infer that transcript attenuation depends on both heteromer formation, and particular cis-element sequences, but how the combination of the two promotes transcriptional termination is unclear. It is also unclear why transcript attenuation is used to inhibit expression, instead of the more commonly observed mechanism of promoter inhibition. At least in this case, the attenuation works in the context of long operon to decrease gene expression in a manner proportional to distance from the binding site. Thus, perhaps partial expression of genes early in the operon is somehow beneficial (Irnov 2010), or there may be timing benefits of targeting a longer window of elongation rather than the instantaneous event of promoter initiation (Belogurov 2015). Finally, we note that activation by SwrA•DegU can override SinR•SlrR during swarming motility, SinR•SlrR dampening can override SwrA•DegU during biofilm formation. Thus expression may be fine-tuned by differential regulation of two processes that act in opposition.

Ultimately, we generate a comprehensive molecular model for both population heterogeneity and the transition from motility to biofilm formation (**Fig 1A**). In conditions where SwrA is either mutated or otherwise low in cytoplasm, the SinR•SlrR heteromer increases the frequency of non-motile chains, and chaining cells have been thought to be a precursor to biofilm formation (Branda 2001; Vlamakis 2008; Chai 2008; Lopez 2009). We also find that hyperactivation of the SinR•SlrR heteromer can override motility gene expression even in the presence of SwrA, as in the artificial case where an extra copy of the *slrA* gene is provided. The *slrA* extra copy condition likely resembles the situation when biofilm formation is activated, where SinR repression fails and the *eps* operon is expressed to produce the EPS polysaccharide component that promotes cohesion (Kearns 2005). As a consequence, EpsE, a protein encoded within the *eps* operon, interacts with the flagellum to rapidly arrest rotation (Blair 2008; Guttenplan 2010; Subramanian 2019), and the SinR•SlrR heteromer to attenuates the *fla/che* operon transcript so that motility-inhibited biofilm cells grow without synthesizing new flagella. Together, a complex system of functional and transcriptional inhibitors operate at fast and slow timescales, to promote and stabilize biofilm development respectively.

## MATERIALS AND METHODS

### Strain and growth conditions

*B. subtilis* strains were grown in lysogeny broth (LB) (10 g tryptone, 5 g yeast extract, 5 g NaCl per liter) broth or on LB plates fortified with 1.5 % Bacto agar at 37°C. The following antibiotic concentrations were used when necessary: ampicillin 100 μg/ml (*amp*), kanamycin 5 μg/ml (*kan*), chloramphenicol 5 μg/ml (*cm*), spectinomycin 100μg/ml (*spec*), tetracycline 10 μg/ml (*tet*), and erythromycin 1μg/ml plus lincomycin 25μg/ml (*mls*).

### Strain construction

*B. subtilis* chromosomal DNA from indicated strains was used to amplify all PCR products. All constructs were transformed into the naturally competent DK1042 that carries a *comI^Q12L^* mutation in the 3610 ancestral strain. SPP1 phage lysate of strains carrying constructs with selectable markers were prepared and transduced into desired genetic backgrounds using generalized transduction.

#### SPP1 phage transduction

Donor *Bacillus subtilis* strains were grown in TY broth (LB broth supplemented with 10 mM MgSO_4_ and 100 µM MnSO_4_). Serial dilutions of SPP1 phage stock were added to 0.2 ml of dense culture (OD600-0.6-1.0) and statically incubated at 37°C for 15 mins. 3 ml of molten TY soft agar (TY supplemented with 0.5% agar) was added to each mixture, poured on top of fresh TY agar plates (TY supplemented with 1.5% agar) and incubated at 37°C overnight. The top agar a plate containing near confluent plaques was scraped and collected in a 15ml conical tube, vortexed and centrifuged at *5,000Xg* for 10 mins. The supernatant that contained phage particles was passed through a 0.45 µm syringe filter to eliminate any bacterial contamination and stored at 4 °C. Recipient strains were grown to OD600-0.6-1.0 in TY broth at 37°C and one ml of cells were mixed with 25µl of SPP1 phage stock from the donor. 9ml of TY broth was added to the mixture and the mixture was incubated at room temperature for 30 mins with gentle shaking on a rocker. The mixture was centrifuged at *5000Xg* for 10 mins, supernatant was discarded, pellet was resuspended in the remaining volume and the 100µl of the cell suspension was plated on LB plates fortified with 1.5% agar, supplemented with the appropriate antibiotics and 10mM sodium citrate. The plates were incubated at 37°C overnight. All strains used in this study are listed in **Table 1**. All primers used to build strains for this study are listed in **Table S8** and all plasmids are listed in **Table S9**.

**TABLE 1:**
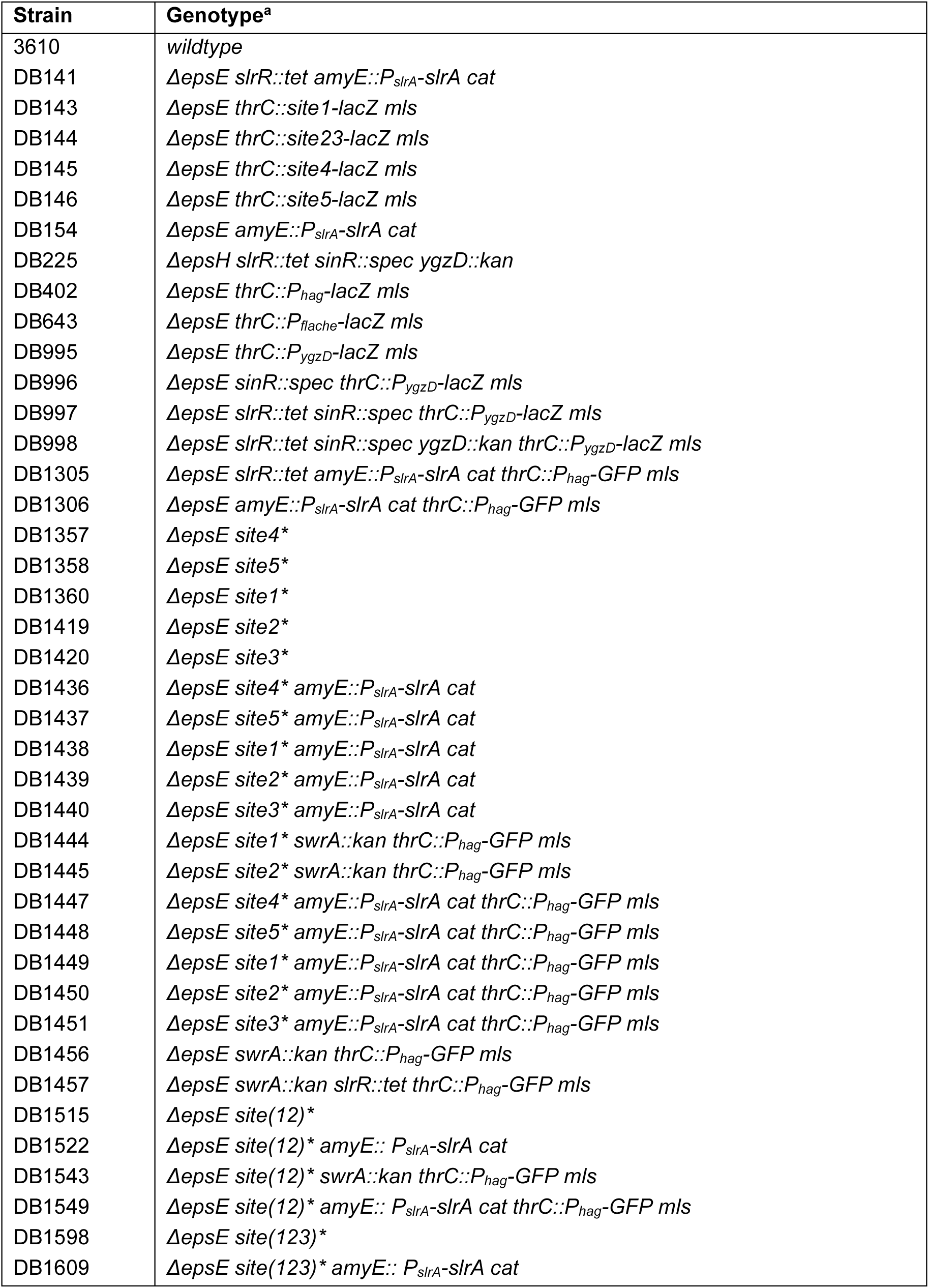
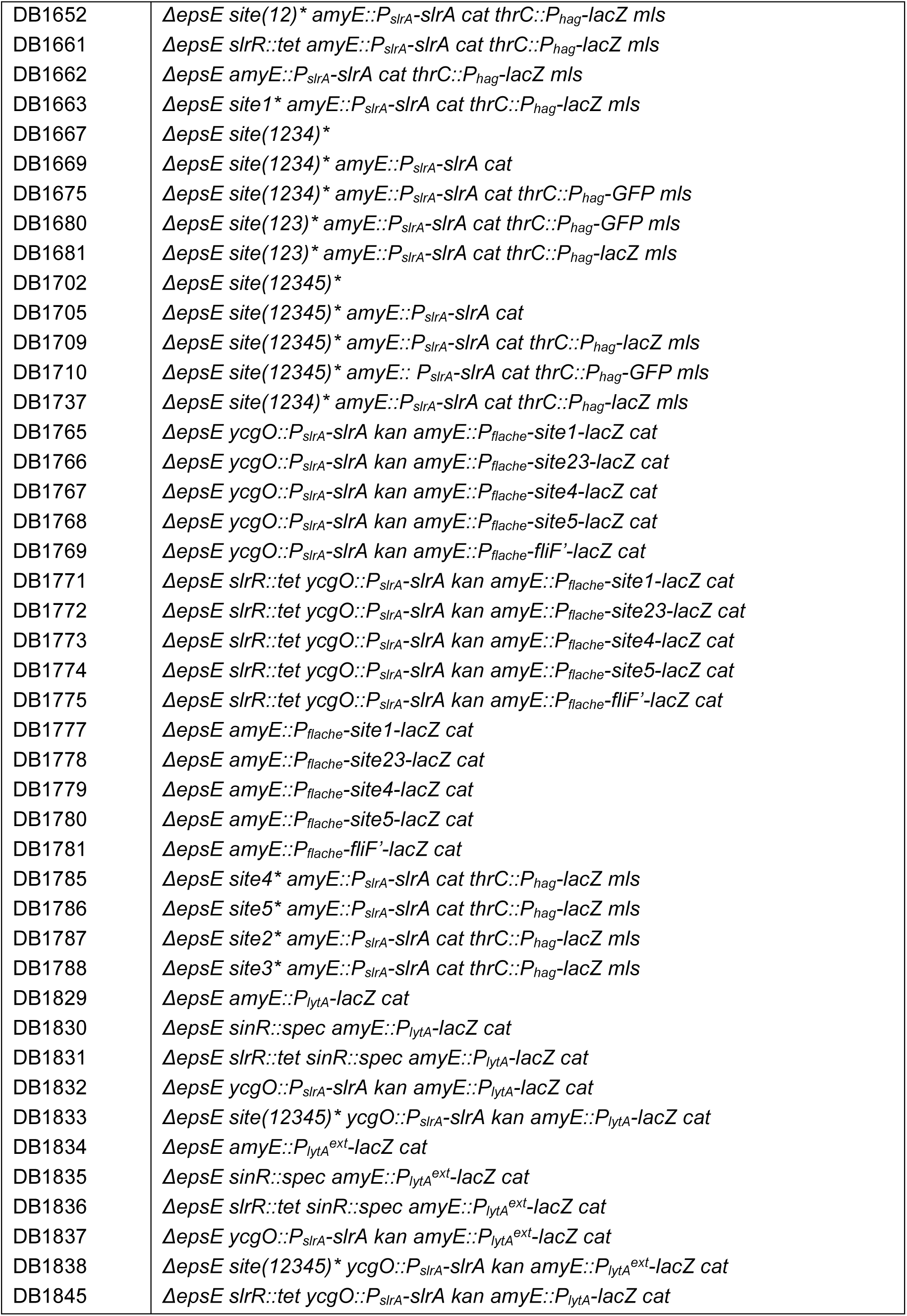
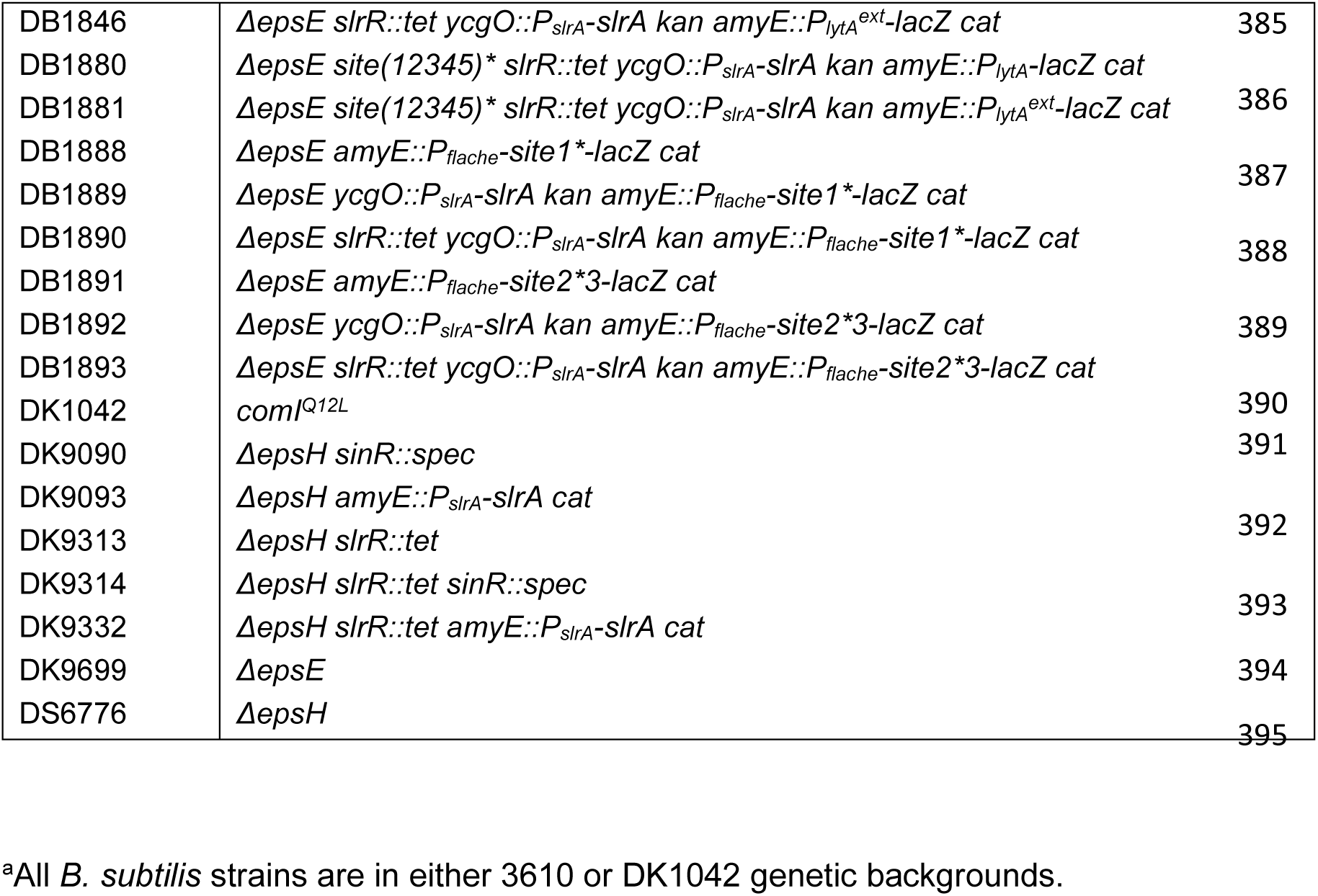
Strains.

#### Transcriptional reporter constructs

##### pAM58, 59, 60, 61, 81, 109

DK1042 chromosomal DNA was used to amplify regions using primer pairs 7928/7929, 7930/7931, 7926/7927, 7924/7925, 8139/8140, 8141/8142 and ligated into the EcoRI/BamHI sites of pDG1663 containing the *lacZ* gene and the gene for *mls* resistance between arms of *thrC* to generate *pAM58, 59, 60, 61, 81* and *109* respectively. These were separately transformed into DK1042 and integration at the *thrC* locus was confirmed by the ability of the mutant to grown on *mls* and the inability of mutant to grow in defined media in the absence of threonine.

##### pAM112

DK1042 chromosomal DNA was used to amplify approximately 500 bp upstream of the *flache* ribosomal binding site using primers 8008/8423 and ligated into the EcoRI/ BamHI sites of pDG268 containing the *lacZ* gene and the *cat* gene for chloramphenicol resistance between arms of *amyE* to generate *pAM112*.

##### pAM113, 114, 115, 116, 117

DK1042 chromosomal DNA was used to amplify ∼500bp regions surrounding *sites 1, 23, 4, 5* and a region within *fliF* using primer pairs 8424/8425, 8426/8427, 8428/8429, 8430/8431 and 8432/8433 and ligated into the NheI/BamHI sites of *pAM112* to generate *pAM113, 114, 115, 116* and *117* respectively. These plasmids were separately transformed into DK1042 and chromosomal integration into the *amyE* site was confirmed by resistance of the transformants to chloramphenicol and their inability to digest sucrose when grown on LB media supplemented with sucrose.

##### pAM118

Chromosomal DNA from *B.subtilis* strain DB1360 was used as a template to amplify 500bp region surround *site1\** mutation using primer pairs 8424/8425 and ligated into the NheI/BamHI sites of *pAM112* to generate *pAM118*. *pAM118* was transformed into DK1042 and the transformants were confirmed as mentioned above.

##### pAM119

Chromosomal DNA from *B.subtilis* strain DB1419 was used as a template to amplify 500bp region surround *site2\** mutation using primer pairs 8426/8427 and ligated into the NheI/BamHI sites of *pAM112* to generate *pAM119*. *pAM119* was transformed into DK1042 and the transformants were confirmed as mentioned above.

#### *slrA* complementation construct

##### pAM103

DK1042 chromosomal DNA was used to amplify approximately 500 bp upstream of the *slrA* ribosomal binding site using primers 8337/8338 and ligated into the EcoRI/BamHI sites of *pKM087* to generate *pAM103*. *pAM103* was transformed into DK1042 and integration into the chromosome was confirmed by the presence of a rugose colony morphology because of the presence of an extra copy of *slrA*.

#### Native site mutants

##### pDP620

DK1042 chromosomal DNA was used to amplify ∼1000bp flanking fragments surrounding *site1* using primer pairs 8273/8274 and 8275/8276 that contained the *site1\** mutation. *pminiMAD3* was linearized by digesting with SmaI and the two flanking fragments were assembled into *pminiMAD3* by Gibson assembly.

##### pDP621

DK1042 chromosomal DNA was used to amplify ∼1000bp flanking fragments surrounding *site23* using primer pairs 8277/8278 and 8279/8280 that contained the *site2\** mutation. *pminiMAD3* was linearized by digesting with SmaI and the two flanking fragments were assembled into *pminiMAD3* by Gibson assembly.

##### pDP622

DK1042 chromosomal DNA was used to amplify ∼1000bp flanking fragments surrounding *site23* using primer pairs 8281/8282 and 8283/8284 that contained the *site3\** mutation. *pminiMAD3* was linearized by digesting with SmaI and the two flanking fragments were assembled into *pminiMAD3* by Gibson assembly.

##### pDP623

DK1042 chromosomal DNA was used to amplify ∼1000bp flanking fragments surrounding *site4* using primer pairs 8285/8286 and 8287/8288 that contained the *site4\** mutation. *pminiMAD3* was linearized by digesting with SmaI and the two flanking fragments were assembled into *pminiMAD3* by Gibson assembly.

##### pDP624

DK1042 chromosomal DNA was used to amplify ∼1000bp flanking fragments surrounding *site5* using primer pairs 8289/8290 and 8291/8292 that contained the *site5\** mutation. *pminiMAD3* was linearized by digesting with SmaI and the two flanking fragments were assembled into *pminiMAD3* by Gibson assembly.

##### pDP638

Chromosomal DNA from *B.subtilis* strain DB1419 was used to amplify ∼1000bp flanking fragments using primer pairs 8281/8397 and 8398/8284 that contained both *site2\** and *site3\** mutations. *pminiMAD3* was linearized by digesting with SmaI and the two flanking fragments were assembled into *pminiMAD3* by Gibson assembly.

The plasmids were passaged individually through *recA+ E. coli* strain TG1, transformed into DK1042 and plated at restrictive temperature for plasmid replication (37°C) on LB agar supplemented with *spec* to select for transformants with single crossover plasmid integration. Plasmid eviction was ensured by growing the strains for 14 hours at a permissive temperature for plasmid replication (22°C) in the absence of *spec* selection. Cells were serially diluted, plated on LB agar plates in the absence of *spec* and individual colonies were replica patched on LB agar plates with and without *spec* to identify *spec* sensitive colonies that have successfully evicted the plasmid. Chromosomal DNA was isolated form the colonies that had excised the plasmid and allelic replacement was confirmed by sequencing.

### Swarm expansion assay

1 mL of mid-log phase cells (OD_600_ 0.3-1.0) grown at 37°C in LB were harvested and resuspended to and OD_600 of_ 10 in pH 8.0 PBS (137 mM NaCl, 2.7 mM KCl, 10 mM Na_2_HPO_4_, and 2 mM KH_2_PO_4_) containing 0.5% India ink (Higgins). Freshly prepared LB plates fortified with 0.7% bacto agar (25 mL per plate) was dried for 10 min in a laminar flow hood, centrally inoculated with 10 μL of the cell suspension, dried for another 10 min, and incubated at 37 °C. The India ink demarks the origin of the colony and the swarm radius was measured relative to the origin every 30 min. For consistency, an axis was drawn on the back of the plate and swarm radii measurements were taken along this transect.

### β-galactosidase assay

*B. subtilis* strains were grown in LB broth at 37°C with constant rotation to OD_600_ 0.7-1.0. One mL of cells was harvested by centrifugation and resuspended in 1 mL of Z-buffer (40 mM NaH_2_PO_4_, 60 mM Na_2_HPO_4_, 1 mM MgSO_4_, 10 mM KCl and 38 mM β-mercaptoethanol). To each sample, lysozyme was added to a final concentration of 0.2 mg/mL and incubated at 30°C for 15 minutes. Each sample was diluted appropriately in 500 μl of Z-buffer and the reaction was started with 100 µl of start buffer (4 mg/ml 2-nitrophenyl β-D-galactopyranoside (ONPG) in Z-buffer) and stopped with 250 µl 1 M Na_2_CO_3_. The OD_420_ of the reaction mixtures were recorded and the β-galactosidase specific activity was calculated according to the equation: (OD_420_/time x OD_600_)] x dilution factor x 1000.

### Chromatin Immunoprecipitation Sequencing (ChIP-Seq)

*Bacillus subtilis* cultures were grown to an OD_600_ of 1.0 at 37°C with constant rotation. 20 mL of cells were cross-linked for 30 minutes at room temperature using 3% formaldehyde (Sigma), quenched with 125 mM glycine, washed with PBS, and then lysed. DNA was sheared to an average fragment size of ∼200 bp using Qsonica sonicator (Q8000R), and then incubated overnight at 4°C with α-SinR (Kearns, 2005). Immunoprecipitation was performed using Protein A Magnetic Sepharose beads (Cytiva #45002511), washed, and DNA was eluted in TES (50mM Tris pH8, 10mM EDTA and 1% SDS). Crosslinks were reversed overnight at 65°C. DNA samples were treated with a final concentration of 0.2 mg/ml RNaseA (Promega #A7973) and 0.2 mg/ml Proteinase K (NEB #P8107S) respectively, and subsequently extracted using phenol/chloroform/isoamyl (25:24:1). DNA samples were then used for library preparation using NEBNext UltraII DNA library prep kit (NEB #E7645L). Paired end sequencing of the libraries was performed using NextSeq 500 platform and at least 3 million paired-end reads were obtained for each sample. Two or three biological replicates were sequenced for each sample.

### Whole genome sequencing (WGS)

*Bacillus subtilis* cultures were grown to an OD_600_ of 1.0 at 37°C with constant rotation and 5 ml of cells were collected, pelleted and DNA was extracted using Qiagen DNeasy kit (#69504). Sonication of genomic DNA was performed using Qsonica sonicator (Q8000R) and the sonicated DNA was used to prepare libraries using the NEBNext UltraII DNA library prep kit (NEB #E7645L). Paired end sequencing of the libraries was performed using NextSeq 500 platform and at least 3 million paired end reads were obtained for each sample. Data from WGS was used as input for the ChIP.

### Analysis of ChIP-Seq and WGS data

Sequencing reads for both ChIP and WGS were mapped individually to *B. subtilis* 3610 genome (NZ_CP020102.1) (Nye, 2017) using CLC Genomics Workbench software (Qiagen). The enrichment at ribosomal RNA locations were eliminated and the number of reads mapped to each base pair in the genome was exported into a .csv file. Data were normalized to the total number of reads and fold enrichment was calculated as the ratio of number of reads at each genome location in ChIP-Seq and WGS (ChIP/input). Analysis was performed and graphs were plotted in 1-kb bins to show enrichment across the entire genome using custom R-scripts. When required, individual peaks were plotted in 10-bp bins across a 4-kb range centered around the peak summit. Detailed protocols and scripts are available upon request.

### MEME analysis

200-bp sequence surrounding the peak center in was extracted using a custom perl script and a fasta file was created. Sequences were subjected to Multiple Em for Motif Elicitation (MEME) v 5.5.2 using parameters (meme sequences.fa -dna -oc . -nostatus -time 14400 -mod anr -nmotifs 3 -minw 21 -maxw 21 -objfun classic -revcomp -markov_order 0). 21 bp highly enriched motif sequences were extracted and sequence logo generated by MEME is presented in **Fig 2B**.

### Microscopy

For microscopy, 3ml of LB broth was inoculated with a single colony and grown at 37°C. 1ml of culture at OD_600_ 0.5-0.8 was pelleted and resuspended in 30µl 1X PBS buffer supplemented with 5 µg/ml FM 4-64 (Invitrogen #T13320) and incubated at room temperature for 2 min in the dark. The cells were washed once with 1mL of PBS, spun down and resuspended in 30μl of PBS. 5μl of sample was spotted onto flat agarose pads (1% agarose in PBS) on slides and covered with a glass coverslip. Fluorescence microscopy was performed with a Nikon 80i microscope with a phase contrast objective Nikon Plan Apo 100X and an Excite 120 metal halide lamp. FM4-64 was visualized with a C-FL HYQ Texas Red Filter Cube (excitation filter 532-587 nm, barrier filter >590 nm). GFP was visualized using a C-FL HYQ FITC Filter Cube (FITC, excitation filter 460-500 nm, barrier filter 515-550 nm). Images were captured with a Photometrics Coolsnap HQ2 camera in black and white using NIS elements software and subsequently false colored and superimposed using Fiji v 2.1.0 (Schindelin, 2012).

### Structure prediction

Multimer structure prediction of SinR•SinR, SlrR•SlrR, SinR•SlrR and YgzD•YgzD was performed using Alphafold2 (Jumper, 2021). For multimer prediction, amino acid sequence for each protein from *Bacillus subtilis 3610* genome (NZ_CP020102.1) was separated by a colon (:) and prediction was performed using parameters colabfold_batch –num-recycle 20 –amber –templates –model-type alphafold2_multimer_v2. Structures were visualized and shaded using UCSF Chimera v 1.15 (Petterson, 2004).

### Sequence alignment

Amino acid sequence of YgzD , SinR and SlrR protein from *Bacillus subtilis 3610* genome (NZ_CP020102.1) were aligned by Clustal Omega v 1.2.4 using default parameters (Sievers, 2011). Alignment was shaded using Jalview v 2.11.2.7 using a 50% identity threshold (Waterhouse, 2009).

### RNA extraction and analysis

RNA was extracted from *B.subtilis* as described earlier (Cozy, 2012) with slight modifications. *B.subtilis* strains were grown in LB overnight, diluted the next day and grown until they an OD_600_ of ∼1.0. 5ml of culture was flash frozen by adding an equal volume of cold methanol that was pre-frozen at −80°C. The mixture was centrifuged at *5000Xg* for 10 min at 4C, supernatant was discarded, and the pellets were stored at −80°C. Pellets were resuspended in 800µl of hot LETS buffer (10 mM Tris-HCl pH 7.4, 50 mM LiCl, 10 mM EDTA pH 8.0, 1% SDS) preincubated at 75°C. The mixture was added to 650mg acid washed glass beads and 600µl hot acid-saturated phenol pH 4.6 pre-incubated at 75°C. The mixture was vortexed for 3 mins and 600µl of Chloroform was added, vortexed for 30s and centrifuged at 3200g for 10 mins at 4°C. 600µl of the top aqueous layer was added to 800µl of hot phenol-chloroform (5:1, 75°C), vortexed for 3 mins and centrifuged at *3200Xg* for 10 mins at 4°C. The aqueous phase was collected and added to an equal volume of isopropanol, mixed by inversion and left at room temperature for 10 mins. The mixture was centrifuged at 4°C for 25 mins at maximum speed. Supernatant was discarded and the pellet was washed with 1ml ice cold 75% ethanol after which the pellet was dried at room temperature for 10 mins and resuspended in 20µl of nuclease-free water at 55°C for 10 mins. 300ml of TRIzol was added, vortexed for 15s and incubated at room temperature for 5 mins. 60ul of chloroform was added, mixed by inverting the tube for 15s and incubated at room temperature for 2 mins. The mixture was centrifuged at 4°C max speed for 15mins. The aqueous layer was collected and mixed with an equal volume of isopropanol, mixed by inverting and incubated at room temperature for 10 mins. The mixture was centrifuged at max speed at 4°C for 25 mins, supernatant was discarded, and the pellet was washed with 200µl of 75% ethanol. The mixture was centrifuged at max speed, ethanol was discarded, and the pellet was air dried at room temperature for 10 mins. Pellet was finally resuspended in 20µl in nuclease free water at 55°C for 10 mins. RNA was treated with RNase free DNase I at 37°C for 30 min (Invitrogen AM2222) according to manufacturer’s instructions. RNA was reextracted the same way as before starting at the TRIzol set. rRNA depletion and library preparation was performed by Indiana University Center for Genomics and Bioinformatics. Paired end sequencing of the libraries was performed using NextSeq 550 platform and at least 5 million paired end reads were obtained for each sample. Reads were mapped against NCIB 3610 genome (NZ_CP020102.1) (Nye, 2017) and TPM (Transcript per kilobase million) were calculated using CLC genome browser. TPM values are presented in **Table S10**.

## DATA AVAILABILITY

ChIP-Seq and RNA-Seq data have been submitted to the Gene Expression Omnibus and are available under accession numbers GSE285064 and GSE285065 respectively. Protocols and scripts used in this study are available upon request.

## ACKNOWLEDGEMENTS

We thank members of the Kearns lab for helpful discussions and members of the Wang lab for assistance with ChIP-seq. Alphafold was performed using IU carbonate, a part of Indiana University Pervasive Technology Institute that is supported by Lilly Endowment, Inc. We thank Indiana University Center for Genomics and Bioinformatics for their help with next-generation sequencing.

## AUTHOR CONTRIBUTIONS

AM: Conceptualization, Data Curation, Formal Analysis, Investigation, Validation, Visualization, Writing – Draft Preparation

AJ: Investigation, Data Curation

WX: Funding Acquisition, Methodology, Resources, Supervision, Writing – Review and Editing DBK: Conceptualization, Formal Analysis, Investigation, Project Administration, Resources, Supervision, Writing – Draft Preparation

